# Population genetics, phylogeography and gene flow of mainland and island vampire bat (*Desmodus rotundus)* populations: an investigation into mainland-island bat movement

**DOI:** 10.1101/2024.01.29.577751

**Authors:** Janine F.R. Seetahal, Daniel G. Streicker, Peter Beerli, Nikita Sahadeo, Philippe Lemey, Manuel J. Sanchez-Vazquez, Alice Broos, Laura Bergner, Vernie Ramkissoon, Ron Mahabir, Praimnauth Tihul, Dane Hartley, Astrid Van Sauers, Gianna Karg, Ryan S. Mohammed, Roman Biek, Christopher A.L. Oura, Christine V.F. Carrington

## Abstract

Movement of animals and plants from mainland populations contributes to the genetic diversity and viability of geographically isolated island biota, but also carries risks of pathogen introductions. The bat fauna of the island of Trinidad reflects species diversity on the neighbouring South American mainland and includes the common vampire bat (*Desmodus rotundus)*. We determined relationships between Trinidad and mainland vampire bat populations and the extent of mainland-island movement by comparing the genetic structure (nuclear and mitochondrial) and morphology of the Trinidadian *D. rotundus* population to populations in neighbouring regions of the South American mainland and estimating evolutionary histories and patterns of gene flow.

Results indicate that Trinidadian *D. rotundus* are genetically and morphologically distinct from mainland populations, although limited unidirectional male-biased mainland to island gene flow occurs at an estimated rate of 3.3 migrants per year. Two geographically-defined *Desmodus* cytochrome *b* clades were identified within Trinidad (i.e., one restricted to the South-Western peninsula which grouped with Venezuelan sequences and the other found throughout the rest of the island which grouped with sequences from Suriname and Guyana) suggesting long-standing female philopatry. The geographic distribution of these clades mirrors that of two previously identified geographically defined rabies virus (RABV) lineages introduced to Trinidad from the mainland. This finding suggests that bat dispersals and RABV introductions occur via both the northern and south-western island peninsulas, with subsequent male-driven intra-island viral spread enabled by bat nuclear homogeneity of these populations. These study findings will contribute to the development of contemporary evidence-based vampire bat population control and rabies prevention programs within island populations.

## Introduction

Bats are found on all continents except Antarctica [1] with 1,447 species recorded to date [2]. Species richness is greater in the tropics [3] with the forests of South America harbouring greater bat species richness than any other area in the world [4]. As volant mammals, relatively few barriers to their dispersal exist so high levels of gene flow between geographically distant bat populations may be anticipated [5–7]. The twin-island Republic of Trinidad and Tobago is the southernmost country in the Caribbean archipelago located just off the eastern coast of South America, with only 17 km and 12 km between Venezuela and Trinidad’s northwestern and southwestern peninsulas respectively. This proximity and the fact that the landmass of Trinidad was once contiguous with that of the mainland influences the island’s bat fauna which is reflective of the neighbouring continent, with 68 species currently documented, including two species of vampire bats, *Desmodus rotundus* (common vampire bat) and *Diaemus youngi* (white-winged vampire bat) [8, 9].

Vampire bat-transmitted rabies in humans was recognized in the early twentieth century on the island of Trinidad [10], and Trinidad and Tobago subsequently became the first country to establish a national vampire bat control program targeting *D. rotundus* on the island of Trinidad [11, 12]. Human encroachment into bat territories after colonization and the introduction of livestock - a readily accessible food supply – was thought to have resulted in rapid expansion of the vampire bat population [12, 13]. The extent of this expansion was in part due to the plasticity of the species to rapidly adapt to the disruption of their natural habitat [9, 12, 14]. *D. rotundus* colonies usually consist of between 20 and 100 bats, with bats sometimes found occupying niches within shared roosts with other bat species in a wide variety of places (e.g. hollow trees, culverts, caves) [15]. Bat-transmitted rabies cases (animals and humans) have to date been restricted to Trinidad. Tobago (which lies 42 km from the northeast coast of Trinidad), has remained free of vampire bats [8] and vampire bat-transmitted rabies [9].

Islands generally form geographically discrete isolated habitats [16–18] and as a result insular species tend to have lower genetic diversity and population sizes than mainland populations due to several related factors (e.g., inbreeding [19–22], founder effects and genetic drift [21–23]). Therefore, gene flow between island and mainland populations may be essential for evolutionary diversification of island species to prevent loss of genetic diversity and species extinction [24]. The spatial isolation of islands also presents a physical oceanic barrier to the introduction of viruses and influences the transmission dynamics of these infectious agents [25]. In this regard, animal dispersal can be a key determinant for the translocation of pathogens over oceanic ecological barriers [26, 27].

Previous phylogeographic work on Trinidadian rabies viruses (RABVs) provided evidence for at least three separate introductions of RABV lineages (*D. rotundus* variant) from the South American mainland that arose around 1972, 1989 and 2004, with each introduction followed by periods of *in situ* spread and evolution [28]. These introductions were proposed to have occurred via infected bats flying across the short distance from the South American mainland to Trinidad [28]. *Desmodus rotundus* have been reported to fly up to 20 km from roost to feeding site in a single night [14] which would allow them to cross the short distance from the South American mainland to Trinidad to feed and return, or travel further inland on the island. The existence and extent to which such mainland-island movement occurs amongst these *D. rotundus* populations is currently unknown but such movement would be expected to be a primary determinant of the frequency of RABV introductions. For example, one possibility is that mainland-island movement is frequent and the episodic nature of rabies outbreaks in Trinidad reflects episodic viral circulation in mainland source populations. Alternately, mainland-island movement may be rare, with a more or less constant presence of rabies on the mainland. In the former scenario, RABV circulation in dispersing populations would be a significant predictor of RABV introductions into Trinidad whereas in the latter case, determinants of mainland to island bat dispersal would be the main driver of RABV outbreaks and critical to predicting these events.

This study aimed to compare the genetic structure of the Trinidadian *D. rotundus* bat population to populations from nearby regions on the South American mainland and to estimate patterns and rates of *D. rotundus* gene flow between the mainland and Trinidad. Since mammalian dispersal is usually sex-biased and has been demonstrated in several bat species [29–34] including *D. rotundus* [35], we evaluated whether mainland-island and intra-island dispersal events were dominated by male bat movement by comparing genetic markers with differing modes of inheritance i.e. microsatellites and the cytochrome b (*cytb)* gene [36].

Microsatellites are highly polymorphic bi-parentally transmitted co-dominant nuclear markers [37] which are especially useful for analysis of demographic history and fine-scale population structure [38, 39]. The *cytb* gene is a maternally inherited mitochondrial gene [40] widely used in mammalian evolutionary studies, particularly with several neotropical bat species including the *Ardops* sp. [41], *Brachyphylla* sp. [41], *Lonchophyllini* sp. [42], *Artibeus* sp. [41, 43, 44], *Glossphaga* sp. [45], *Uroderma* sp. [46] and *Desmodus* sp. [43] including the vampire bat [43]. Finally, since *D. rotundus* subspeciation was more recently suggested within its continental range due to genetic and morphological variations [47, 48], morphometric features were compared to evaluate the extent of morphological divergence between island and mainland populations.

## Materials and Methods

### Vampire bat trapping, sample collection and DNA extraction

Field trapping of common vampire bats (*D. rotundus*) was conducted in Trinidad (February 2012 -July 2017), Suriname (April 2016) and Guyana (December 2015) in accordance with established guidelines [49]. Bats were captured at ground level, at dusk and during the night at foraging grounds using mist nets and at roosting sites using hand-held nets and a harp trap. Foraging grounds were identified based on reports of bat biting attacks on livestock and were included to prevent familial over-representation. Bats were transported to the laboratory in individual mesh bags and / or small animal cages for processing and sample collection.

Field identification keys were used for morphological species identification [50], and biometrics (i.e. forearm length, weight, head and body length), age (juvenile / adult), sex and reproductive status were recorded. All captured bats used in the study were apparently healthy adults and were each sampled once. Bats were humanely euthanized by anaesthetic overdose using isoflurane and organ tissues were harvested by dissection and stored in-house for other parallel studies. Blood was collected by cardiac puncture and sera was separated by centrifugation and stored until use. All samples and serum were stored at −80⁰C until further processing.

Non-contemporary supplemental samples were sourced from museum collections of voucher bat specimens including: The American Museum of Natural History (AMNH), New York, USA; The Biodiversity Institute and Natural History Museum of Kansas University, Kansas, USA; The Royal Ontario Museum (ROM), Ontario, Canada; The Institut Pasteur de la Guyane, Cayenne, French Guiana and from the Université de Montpellier, France.

Wing membrane (plagiopatagium) punch biopsies approximately 4mm in diameter were collected from the specimens and stored in RNAlater at −80°C until further processing. DNA was extracted from ∼25 mg of wing membrane tissue using the Qiagen® DNeasy Blood and Tissue Kits (Qiagen® Valencia, CA, USA) according to the manufacturer’s instructions. DNA quantity was estimated using a Nanodrop 1000 (Thermo Fisher Scientific Products) and aliquots were stored at −80°C until further processing.

In total, DNA was obtained from 194 bats representing 31 localities in Trinidad (114 bats from 18 sites), Guyana (14 bats from 3 sites), Suriname (19 bats from 2 sites), Venezuela (24 bats from 8 sites) and French Guiana (23 bats from 2 sites). The locations sampled are shown in **Figure 1** and full details of all bats and samples used in the study are provided in **S1 Table**.

**Figure 1:**
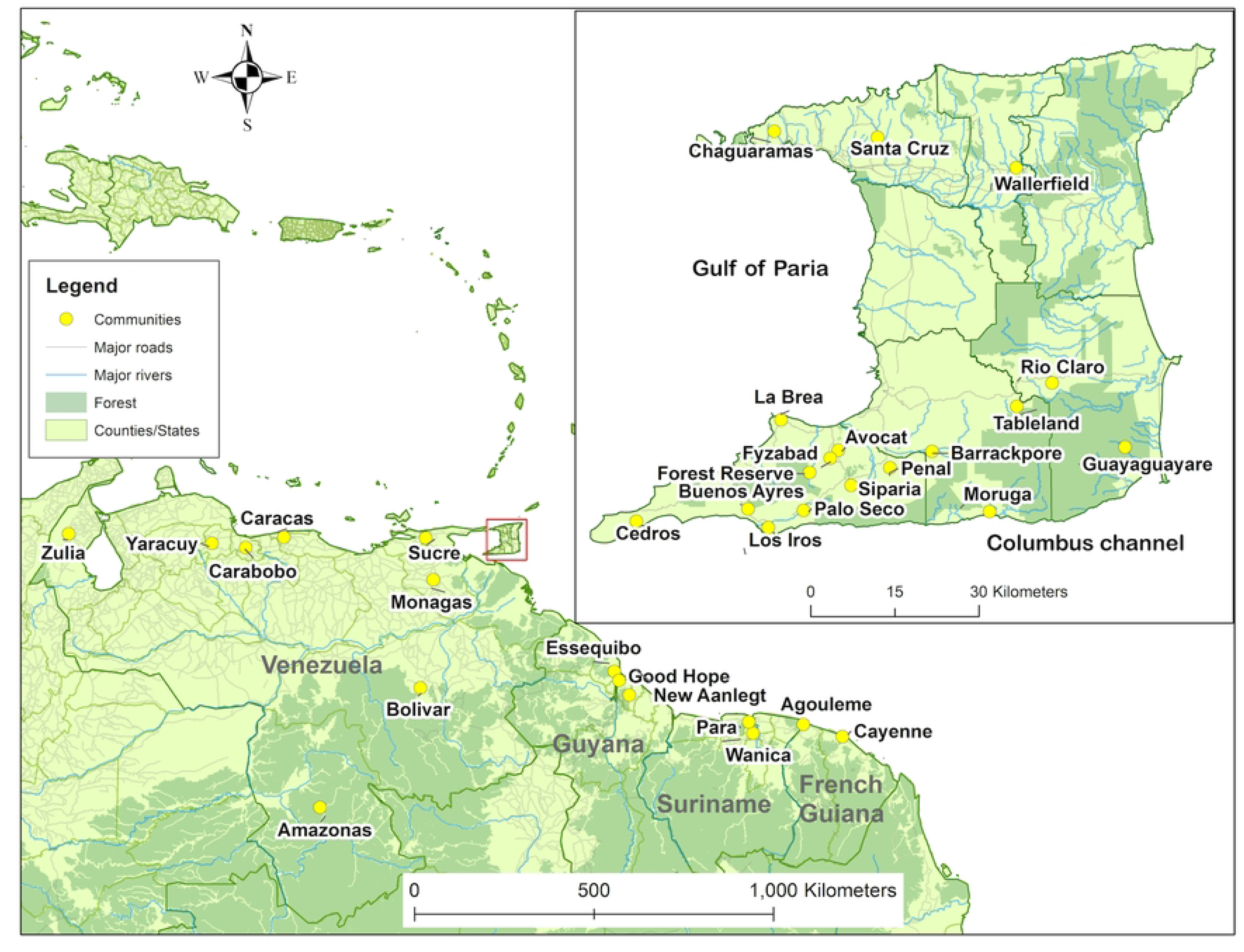
*D. rotundus* sampling locations in Trinidad and South America (Suriname, Guyana Venezuela and French Guiana) for this study.

### Ethical approval and licences

All protocols were approved by The Ethics Committee of the Faculty of Medical Sciences, The University of the West Indies (UWI), St. Augustine Campus (IRB Nos. CEC-19/05/2011-01 and CEC-14/02/2014-02). Field trapping in Trinidad was carried out under annually issued special game licenses from the Wildlife Section, Forestry Division, Ministry of Agriculture, Land and Fisheries, Trinidad and Tobago and were coordinated with the vampire bat population control activities of the Anti-Rabies Unit (ARU) of the Ministry of Agriculture, Land and Fisheries as far as possible. Field collections in Suriname and Guyana were conducted under the authority of the Ministry of Agriculture, Animal Husbandry and Fisheries (LVV) in Suriname and the Guyana Livestock Development Authority (GLDA) in Guyana respectively. In Guyana, trapping was carried out in collaboration with the vampire bat species reduction program of the GLDA. Vampire bats were not protected by law in the countries where field sampling was conducted.

### Mitochondrial cytochrome *b (cytb)* gene amplification and sequencing

*Cytb* was amplified from *D. rotundus* DNA samples (**S2 Table**) using the HotStarTaq® Master Mix Kit (Qiagen® Valencia, CA, USA) and primers described by Martins *et al* [51] at 0.2 µM each, 200 µM each dNTP, 1.5 mM MgCl_2_ and 2.5 units of HotStarTaq® DNA polymerase in a final reaction mixture volume of 50 µl. PCR conditions were based on previously described protocols [51, 52] as follows: 94°C for 15 min; 35 cycles of (94°C for 30 sec, 45°C for 45 sec, 72°C for 70 sec); 72°C for 10 min. Amplicons (1,140bp) were sent to Macrogen Inc. (Seoul, South Korea) for purification and Sanger sequencing with the aforementioned primers. Sequences derived were submitted to Genbank (see **S2 Table** for accession numbers).

### Nuclear microsatellite loci amplification and genotyping

Two multiplex PCRs were used to amplify nine dinucleotide microsatellite loci using primer panels developed by Piaggio *et al.* [53] and the Qiagen® Multiplex PCR kit according to manufacturer’s instructions. PCRs were conducted on each bat genomic DNA sample (*n* = 194), under conditions outlined by Streicker *et al.* [52]. DNA genotyping was performed using an Applied Biosystems 3730xl 96-capillary Genetic Analyser at the University of Dundee, DNA Sequencing and Services, Scotland (https://www.dnaseq.co.uk/).

### Mitochondrial *(cytb)* phylogenetic analysis

*D. rotundus cytb* sequences derived during the current study (n = 64) were aligned with 14 previously published *cytb* gene sequences from *D. rotundus* (n = 10), *Diphylla ecaudata* (n = 2)*, Phyllostomus hastatus* (n =1) and *Phylloderma stenops* (n = 1) using the ClustalW alignment tool within Geneious version 7.1.9 (Biomatters Limited). Alignments were manually examined for gaps and errors, adjusted accordingly then trimmed to a common length of 535bp, corresponding to nucleotide position 60 – 595 of the *D. rotundus cytb* gene complete cds (Genbank acc no. FJ155477). The resulting data set was designated Cytb78_535. Accession numbers and other details are provided in **S2 Table**. Pairwise percent nucleotide sequence identities were determined for all sequences in the data set and the range of identities for the *D. rotundu*s sequences were compared between and amongst countries using Geneious version 7.1.9. (Biomatters Limited), see **S3 Table**.

To determine phylogenetic relationships between these *D. rotundu*s sequences, a maximum likelihood (ML) tree was inferred using IQ-TREE [54] under the best-fit nucleotide substitution model (i.e. TPM2+F+G4) selected by ModelFinder [55] with both ultrafast bootstrap (UFBoot) [56] with 1000 bootstrap samples and SH-aLRT test performed. The consensus tree was then visualized and annotated using Figtree (version 1.4.4.). Nodes with SH-aLRT ≥ 80% and UFboot ≥ 95% were considered strongly supported. To facilitate comparison with a wider range of mainland *D. rotundus* sequences (i.e. including Genbank *D. rotundus cyt-b* gene sequences less than 535bp), a second ML Tree was inferred from a data set (i.e. Cytb154_382) comprised of the cyt-b sequences derived in this study (n = 64) and 90 Genbank sequences (88 *D. rotundus,* 2 *D. ecuadata*) trimmed to a common length of 382bp, corresponding to nucleotide position 24 – 406 of the *D. rotundus cyt-b* gene complete cds (Genbank acc. no. FJ155477).

### Analysis of nuclear genetic structure

Genotyping output data was analysed using the fragment analysis plugin [57] available in Geneious (version 7.1.9.). The output was converted to appropriate formats using GenAlEx (version 6.503) [58] for further analysis using various population genetics software including Microchecker [37] to identify genotyping errors due to null alleles, short allele dominance (i.e. large allele dropout) and scoring errors due to stuttering. The program estimated the null allele frequency and adjusted the observed allele and genotype frequencies before further population genetic analysis. Arlequin version 3.5 [59] was also used to calculate allelic frequencies, F statistics, Hardy-Weinberg equilibrium (HWE) and expected and observed heterozygosities.

Bayesian clustering implemented by STRUCTURE (version 2.3.4) [60] was used to assign individuals to distinct subpopulations or genetic clusters assuming admixture and correlated allele frequency. Samples that could not be scored at six or more microsatellite loci were excluded, to form a data set of 182 individuals (9 loci (*n*= 157); 8 loci (*n* = 15); 7 loci (*n* = 3) and 6 loci (*n* = 7)). The model was run for 30 iterations from K=1 to K=8 with 1,000,000 Markov Chain Monte Carlo (MCMC) repetitions and a burn-in of 10,000 steps. Sampling locations were not used as priors to assist with clustering (i.e. locprior was not employed). Results were then interpreted using the web-based applications STRUCTURE Harvester [61] and Clustering Markov Packager Across K (CLUMPAK) (version 1.1) [62], which both use the Evanno method [63] to estimate the optimum number of population clusters. Individuals with population group membership coefficients q ≥ 0.5 were assigned to that specific genetic cluster.

### Analysis of nuclear genetic connectivity and gene flow

A microsatellite data set was created (*n*=194) with the Trinidad samples split into two subgroups corresponding to the Trinidadian mitochondrial lineages in line with the results of the mitochondrial *cytb* gene analysis. Migrate-n version 3.7 [64] was used on this data set to test eleven models of bat movement (**S1 Figure**) amongst the Trinidadian mitochondrial clades and mainland South America to determine if there were independent migration events per subgroup and migration between subgroups. To improve convergence, we used 40 randomly selected individuals from each Trinidadian subpopulation and all individuals from all other locations. Each model was run in two concurrent MCMC replicates per locus, with 20,000,000 generations per replicate, sampling every 1,000 generations and discarding the first 50% as burn-in. We used an exponential prior distribution for Theta (4Ne*mu) and M (m/mu) with bounds of 0 and 200 (mean M=10) for the immigration parameter. All runs employed heating using 1 cold chain and 3 heated chains with heating parameters of 1.5, 3.0 and 1,000,000. Bayes factors were calculated from the marginal likelihoods provided by the Bezier approximation in the Migrate-n output. In line with previous studies [48] generation time was assumed to be one year.

### Statistical analysis of bat morphometric data

Morphological diversity among *D. rotundus* bats from three different countries, i.e. Guyana (*n*=12), Suriname (*n*=19) and Trinidad (*n*=124), were assessed according to body weight (grams) and forearm length (mm). These variables were modelled using a Gaussian generalised linear model (GGLM) according to country together with sex, to account for its potential confounding effect. The goodness of fit metric, Akaike’s Information Criterion (AIC), was used to compare nested models (i.e., with and without sex) and the Wald test was used to calculate the significance (p-value <0.05) of the variables retained in the final model, particularly for that with multiple categories (i.e. country). Residual diagnostic plots were used to detect features of concern in the model and to identify the presence of potential outliers. All the analyses and graphs were performed using the R statistical software environment [65].

## Results

### Phylogenetic analysis of *cytb* gene sequences

The ML tree estimated from the data set Cytb78_535 is shown in Figure 2 together with the geographic distribution of the *D. rotundus* clades identified. All *D. rotundus* sequences grouped together with very strong support (SH a-LRT 100%; UF bootstrap 100%), with two major *D. rotundus* clades (SH a-LRT 96.8%; UF bootstrap 98%), i.e., DR1 containing Trinidad sequences from the southwest peninsula and all of the Venezuelan sequences, and DR2 which contained the remaining Trinidad sequences (from the southeast and north of the island) together with sequences from Suriname, Guyana, Peru, Mexico and Brazil. The Trinidad DR2 sequences clustered separately from other members of the DR2 clade with moderate support (SH a-LRT 88.1%; UF bootstrap 91%), and were more closely related to sequences from Suriname and Guyana, than those from Peru, Mexico and Brazil.

**Figure 2:**
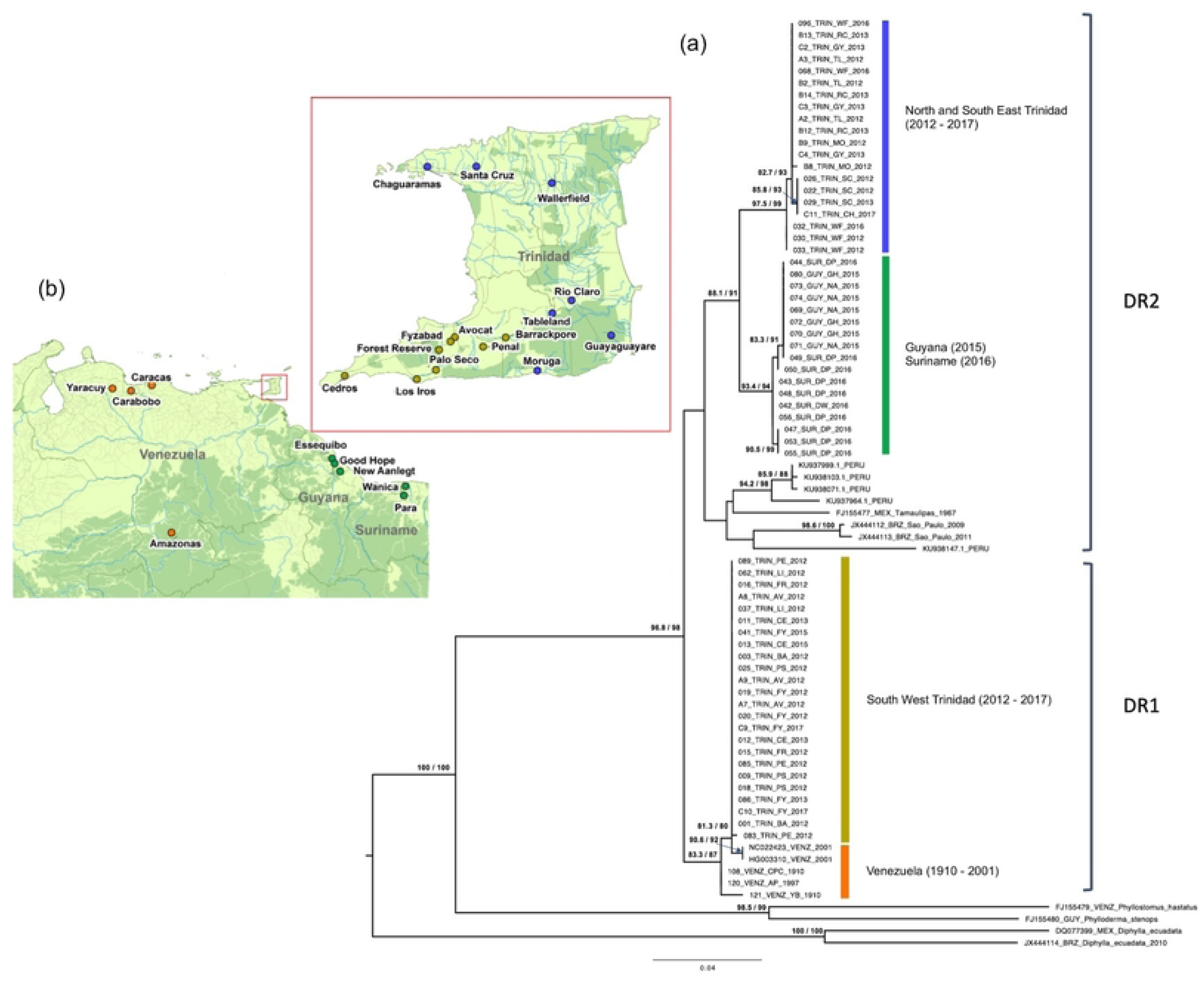
(a) ML tree based on *D. rotundus cyt b* sequences (*n*=78; 535bp) from Trinidad and mainland South American countries together with corresponding sequences from *P. hastatus, P. stenops* and *D. ecuadata*. Taxon labels include sample ID, country, within country location of bat field collection (where available) and year of bat collection, with abbreviation as described in **S1 Table**. SH-aLRT and UFboot ≥ 80% are indicated at selected nodes (SH-aLRT/UFboot). (b) Geographic distribution of *D. rotundus cyt b* sequences belonging to clades DR 1 and DR 2 across sites sampled in this study: Blue circles = Trinidad DR2, olive green = Trinidad DR1, orange circles = Venezuela DR 1 and forest green circles = Guyana and Suriname DR1 sequences.

The topology of the ML tree for the Cytb154_382 data set (**S2 Figure**), which included additional sequences from Latin America was similar, with sequences from this study falling into two major clades (UF bootstrap 77%) corresponding to DR1 and DR2. This larger phylogeny also showed that the DR2 sequences from Trinidad, Venezuela, Guyana and Suriname derived in this study clustered with Brazilian sequences belonging to the previously defined Amazon Forest Clade [48, 51, 66] and were distinct from Brazilian sequences from Atlantic Forest regions [48, 51], as well as those from Peru, Mexico and Central America (Costa Rica, Honduras, El Salvador).

### Genetic diversity: mainland versus island

A total of 84 alleles were detected through microsatellite genotyping of the nine microsatellite loci for island and mainland bats (*n*=194) with the average gene and allelic diversity at sampled locations ranging from 0.15 and 6.6 to 0.69 and 8.3 (**Table 1** and **S4 Table**). There was evidence for null alleles at several loci (**S5 Table**), however the frequencies were generally low (< 0.2) and not expected to significantly bias subsequent analyses [66].

**Table 1:** Overview of gene and allelic variability: averages over nine loci and standard deviation from microsatellite data.

| Population | Gene Diversity | Allelic Diversity | HWE | HE<sub>obs</sub> | HE<sub>exp</sub> | F<sub>IS</sub> | N |
| --- | --- | --- | --- | --- | --- | --- | --- |
| Trinidad | 0.51+/-0.28 | 7.3±3.1 | 4/9 | 0.50±0.31 | 0.53±0.30 | 0.17±0.24 | 228 |
| Suriname | 0.49+/-0.30 | 7.0±2.9 | 6/8 | 0.67±0.28 | 0.69±0.26 | 0.45±0.16 | 38 |
| Guyana | 0.56+/-0.38 | 6.7±1.8 | 6/7 | 0.73±0.11 | 0.80±0.07 | 0.09±0.13 | 28 |
| Venezuela | 0.15+/-0.24 | 6.6±2.5 | 2/9 | 0.39±0.18 | 0.69±0.21 | 0.03±0.17 | 48 |
| French Guiana | 0.69+/-0.39 | 8.3±3.4 | 4/9 | 0.57±0.25 | 0.72±0.29 | 0.20±0.15 | 46 |

HWE: number of loci out of the total polymorphic loci that are in Hardy Weinberg Equilibrium; HE<sub>obs</sub>: observed heterozygosity; HE<sub>exp</sub>: expected heterozygosity; F<sub>IS</sub>=homozygosity index and; N: the number of total allele samples over all loci.

All pairwise comparisons of fixation index (Fst) (i.e. population differentiation) between Trinidad bats and those from mainland countries were significant (p<0.05) (**Table 2**) with the greatest genetic difference noted between the populations sampled from Trinidad and French Guiana (0.103). As shown in **Table 1** and **S3 Figure**, observed and expected heterozygosity (HE_obs_ and HE_exp_) and homozygosity (H_obs_ and H_exp_) were similar for the Trinidad bat population but not for all populations sampled from mainland countries.

**Table 2:** Pairwise genetic distances between populations based on pairwise Fst values from microsatellite data.

| Population | Trinidad | Suriname | Guyana | Venezuela | French Guiana |
| --- | --- | --- | --- | --- | --- |
| Trinidad | 0.000 | - | - | - | - |
| Suriname | 0.0388* | 0.000 | - | - | - |
| Guyana | 0.0463* | 0.0034 | 0.000 | - | - |
| Venezuela | 0.0956* | 0.0085 | 0.0108 | 0.000 | - |
| French Guiana | 0.1030* | 0.0385* | 0.0126 | 0.0239* | 0.000 |

Genetic distances denoted by an asterisk (\*) are significantly different from zero ( $p < 0.05$ )

### Population structure based on microsatellite markers

The female: male ratio for bat specimens used for microsatellite genetic analyses was roughly equal with 84 female bats, 89 male bats and 21 bats unidentified by gender (*n*=194). Bayesian clustering implemented by STRUCTURE [60], using microsatellite data, detected subpopulation structure within the sampled *D. rotundus*, which was best explained by two population clusters with varying levels of admixture for each individual (Figure 3 and **S4 Figure**). Individuals with population group membership coefficients q ≥ 0.5 were assigned to that specific genetic cluster. Most individuals from Trinidad were assigned with high probability to population group 1 and those from the mainland (Guyana, Suriname, Venezuela and French Guiana) were primarily assigned to population group 2. The exceptions were five individuals sampled in Trinidad (Mt1 - Mt5) with atypical admixture, suggesting recent migrant ancestry and one from Guyana (Mg1) and two from Suriname (Ms1 and Ms2) with admixtures in favour of population group 1, which was associated with island bats. Mt1 - Mt5 were mainly male (80%), primarily from collections during earlier years, and were collected from both the northern (Santa Cruz and Wallerfield) and southern (Palo Seco, Buenos Aryes and La Brea) regions of the island (Figure 3).

**Figure 3:**
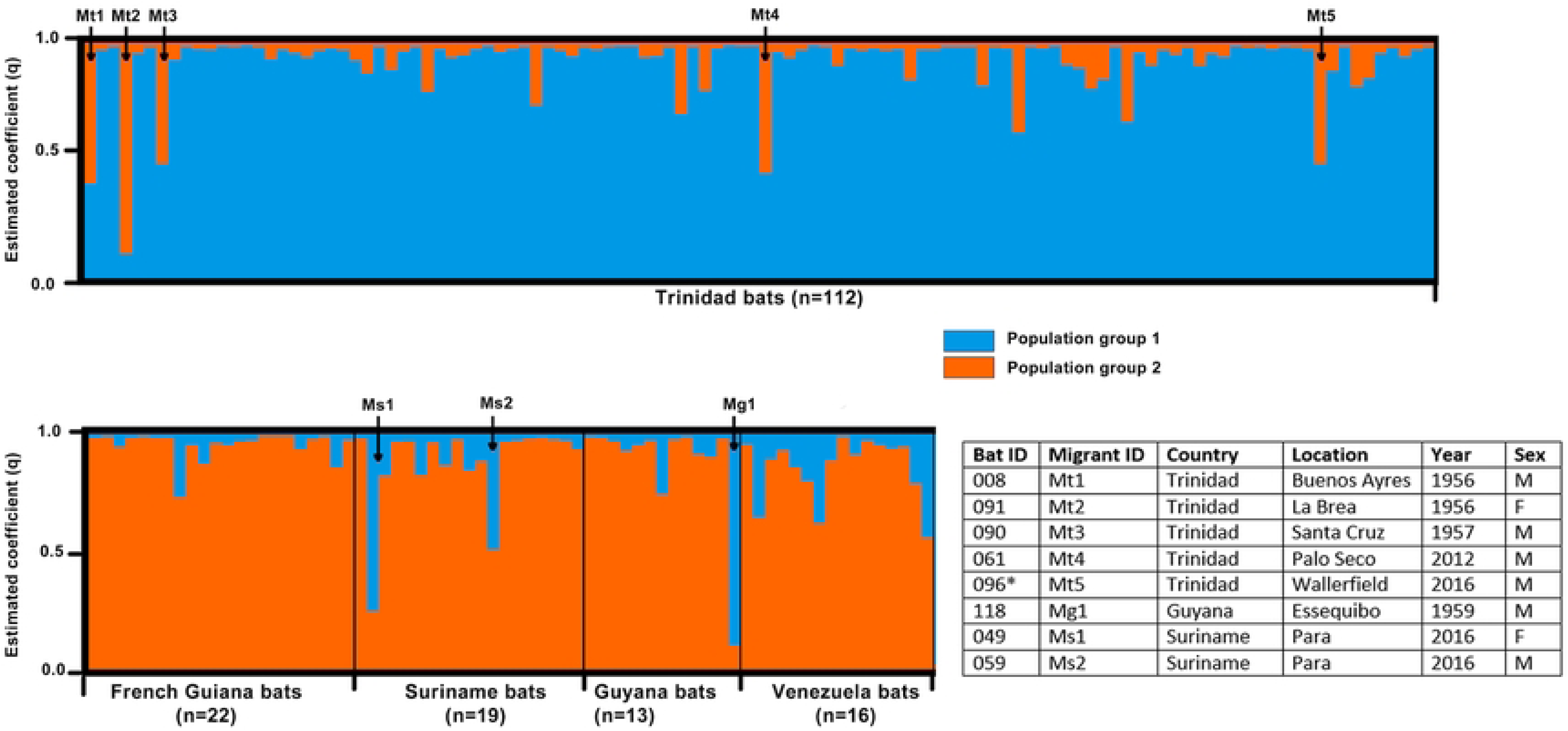
**Structure plots under the assumption of k=2 populations using nine microsatellites (*n*=182 bats)**. Each vertical bar represents the probability of membership assignment of an individual bat to each of the k groups. Black arrows (with Bat ID numbers) indicate putative migrants (atypical population group membership coefficients q≥ 0.5) into the respective countries. The inset table shows the sex, location and date of capture of putative migrant bats. Note: Bat 096* was also included in *cytb* gene analysis (i.e. sequence: 096_TRIN_WF_2016) and belonged to TRIN DR2 *cytb* clade.

### Comparison of models of *D. rotundus* gene flow

Using a model selection approach with the program Migrate-n, different migration models were compared for movement of *Desmodus* bats amongst the two geographically defined Trinidadian *cytb* clades and the South American mainland (**Table 3**). The data best supported the model for one-way bat movement from a panmictic population on the mainland to a panmictic population on Trinidad with no distinction between Trinidadian clades. The mean number of migrants per generation (per year) was estimated to be 3.3 individuals (95% credibility interval = 0.0 to 12.1).

**Table 3:**
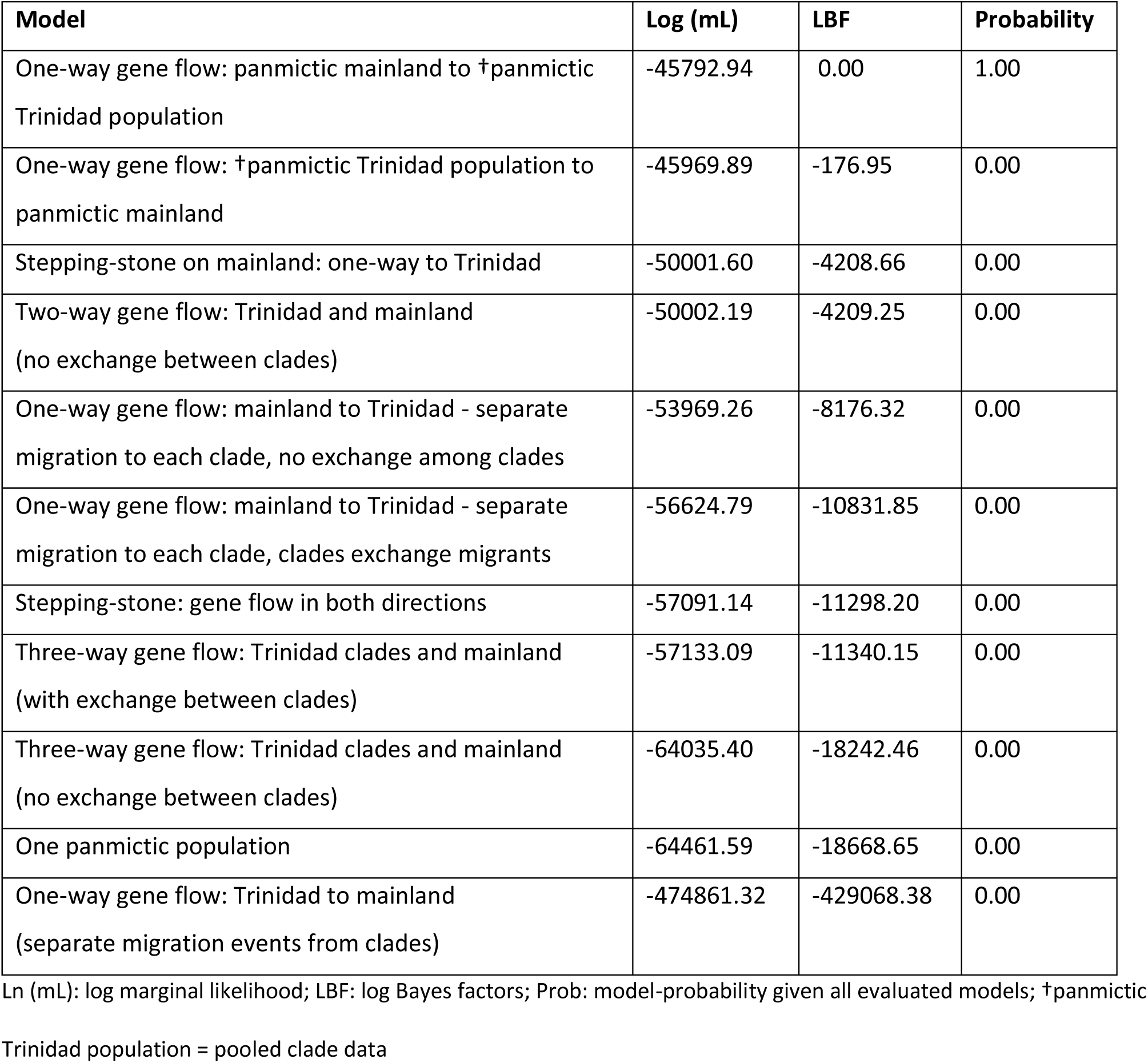
Model comparison for bat movement between Trinidadian clades and the mainland and amongst Trinidadian clades using microsatellite data.

| Model | Log (mL) | LBF | Probability |
| --- | --- | --- | --- |
| One-way gene flow: panmictic mainland to †panmictic Trinidad population | -45792.94 | 0.00 | 1.00 |
| One-way gene flow: †panmictic Trinidad population to panmictic mainland | -45969.89 | -176.95 | 0.00 |
| Stepping-stone on mainland: one-way to Trinidad | -50001.60 | -4208.66 | 0.00 |
| Two-way gene flow: Trinidad and mainland (no exchange between clades) | -50002.19 | -4209.25 | 0.00 |
| One-way gene flow: mainland to Trinidad - separate migration to each clade, no exchange among clades | -53969.26 | -8176.32 | 0.00 |
| One-way gene flow: mainland to Trinidad - separate migration to each clade, clades exchange migrants | -56624.79 | -10831.85 | 0.00 |
| Stepping-stone: gene flow in both directions | -57091.14 | -11298.20 | 0.00 |
| Three-way gene flow: Trinidad clades and mainland (with exchange between clades) | -57133.09 | -11340.15 | 0.00 |
| Three-way gene flow: Trinidad clades and mainland (no exchange between clades) | -64035.40 | -18242.46 | 0.00 |
| One panmictic population | -64461.59 | -18668.65 | 0.00 |
| One-way gene flow: Trinidad to mainland (separate migration events from clades) | -474861.32 | -429068.38 | 0.00 |

Ln (mL): log marginal likelihood; LBF: log Bayes factors; Prob: model-probability given all evaluated models; †panmictic
Trinidad population = pooled clade data

### Morphometric diversity

The mean and standard deviation (SD) for weight and forearm length between contemporary bat populations from Trinidad and the mainland are presented in **Table 4**. Bats from the mainland (Guyana and Suriname) appeared to have greater mean weights and forearm lengths than those from Trinidad; with Guyanese bats demonstrating the largest morphometric characteristics. The results for the GGLMs confirmed these observed differences. Thus, the estimates for the weight model indicated that bats from the mainland were significantly heavier than those from Trinidad (**Table 5**). Likewise, the forearm length model indicated that mainland bats had significantly greater forearm length than Trinidadian bats (**Table 5**).

**Table 4:** Morphometric descriptions of Trinidad, Guyana and Suriname *D. rotundus* populations by weight and forearm length.

| Country | N | Weight (g) |  | Forearm length (mm) |  |
| --- | --- | --- | --- | --- | --- |
|  |  | Mean | S.D. | Mean | S.D. |
| Trinidad | 124 | 24.55 | 3.98 | 56.69 | 2.60 |
| Guyana | 12 | 33.08 | 6.95 | 60.67 | 2.93 |
| Suriname | 19 | 27.11 | 2.73 | 59.58 | 2.43 |

S.D.: standard deviation

**Table 5:** Estimated coefficients for the GGLM for bat body weight and forearm length against country accounting for the effect of sex. N=155.

| Variables | Level | Weight (g) |  | Forearm length (mm) |  |
| --- | --- | --- | --- | --- | --- |
|  |  | Estimate<br>coefficient | 95% CI | Estimate<br>coefficient | 95% CI |
| Sex<br>(baseline = male) | Female | 1.99 | 0.69 - 3.28 | 1.68 | 0.82 - 2.38 |
| Country<br>(baseline = Trinidad) | Guyana | 8.60 | 6.21 - 10.99 | 4.03 | 2.56 - 5.50 |
|  | Suriname | 2.83 | 0.89 - 4.77 | 3.11 | 1.91 - 4.31 |

## Discussion

Geographically distinct animal populations may be morphologically and / or genetically divergent due to selection and genetic drift within separate gene pools. These effects may also vary for different genetic markers which display independent rates of evolution and mutation. In light of the short distance between the island of Trinidad and the South American mainland, low population differentiation of the *D. rotundus* bat populations with free gene flow between localities may have been anticipated, as seen in other areas [67]. However, our results indicate distinct genetic differentiation with limited unidirectional mainland to island gene flow between populations, suggesting the presence of a barrier that hinders dispersal of *D. rotundu*s between Trinidad and the mainland. The Gulf of Paria and Columbus Channel (water bodies between Venezuela and Trinidad) are the most obvious barriers to free movement. However, land barriers within Venezuela (such as deforested or highly urbanised areas) cannot be ruled out since coastal populations in Venezuela were not among those sampled.

In general, analysis of the microsatellite nuclear marker identified two distinct but admixed genetic populations of *Desmodus*, one primarily in Trinidad and the other on the mainland, each including some alleles from the other population. The presence of alleles from these two population groups in both island and mainland bats indicates that although low, some bat movement occurs with successful mating imparting allele transfer [36]. Recent migrant ancestry was inferred for several individual bats from both Trinidad and the South American mainland (see Figure 3), further implying bat movement. Our results suggest that this movement was largely from a panmictic mainland population into Trinidad, and are likely to be primarily male-driven due to the absence of shared (maternally inherited) mitochondrial DNA between mainland and Trinidadian bat populations, which is consistent with female philopatry [68, 69]. Dispersal movement from the mainland to Trinidad was estimated at about 3 individuals per year. However, low level movement in the reverse direction (from island to mainland) cannot be completely ruled out.

In contrast to the nuclear marker which showed no evidence of population subdivision within Trinidad, phylogenetic analysis of *cyt b* distinguished two spatially defined mitochondrial lineages of Trinidadian *Desmodus* sequences (belonging to clades DR1 and DR2). The Trinidadian sequences from the southwest region of the island (DR1 sequences) were more closely related to sequences from Venezuela than to those from other parts of the island (i.e., DR2 sequences), while the latter were more closely related to (but clustered separately from) sequences from Guyana and Suriname. This DR1 pattern suggests more contemporary bat movement between the mainland and the southwest peninsula of the island while the DR2 lineage may be reflective of more historical connectivity between these bat populations and the mainland and island land masses.

Studies that utilised the *cytb* mitochondrial gene to examine *D. rotundus* populations in Costa Rica [51], Peru [52], Uruguay [70], Colombia [71], Mexico [72], French Guiana [51, 72], Brazil [51, 72–73] and Ecuador [74] have shown distinct mitochondrial lineages characterized by Pleistocene vicariance [73] and limited dispersal due to geographical barriers (e.g. the Andes mountains) [52, 74] and or biogeographic landscape features (e.g., forest biomes in Brazil) [51, 73]. However, within the island of Trinidad, outside of the mountain ranges for which the highest elevation is only about 300 m (1,000 ft), there are currently no obvious major geophysical barriers to bat dispersal and therefore, contemporary maintenance of DR1 and DR2 mitochondrial structure (i.e. lack of female gene flow between clades) may likely be due to female philopatry, and ecological and / or anthropogenic factors. For example, habitat fragmentation with discontinuity in resource availability may pose ecological barriers to bat movement [73]. In Trinidad, forest fragmentation is notable between the southwest and southeast forest reserves (see **S5 Figure**), which correlates with the mitochondrial pattern of Trinidadian *Desmodus* population subdivision. Land use data also indicates that within this area the vegetation cover is dominated by hardwood lumber trees, *Tectona grandis* (Teak) and *Peltogyne floribunda* (Purple heart) [75], that would not typically support tree roosting habitats for vampire bats.

The historical structuring of the Trinidadian vampire bat population could also be maintained by anthropogenic environmental alterations associated with human residential settlements (e.g. artificial lighting) which could limit vampire bat foraging and effectively create barriers to mitochondrial gene flow. Finally, the distribution of livestock farms in the southern region of the island (**S5 Figure**), may also contribute to the maintenance of the population structure for the Trinidadian *Desmodus* bats, as these farms may provide reliable food sources and discourage bat movement to other areas [76]. It may also be worthwhile investigating if similar population subdivision occurred for other vertebrate species within these ecodomains, particularly less mobile mammals in which the effect of vicariance might be more pronounced.

Previous studies on the genetic diversity of the common vampire bat have generally focused on their continental range from Central America (e.g., [76–79]) to South America (e.g., [51, 52, 72, 73]), as such, a holistic view of the *Desmodus* population structure has not been conducted along its entire geographic range inclusive of island populations. Some studies have shown greater population subdivision in northern *Desmodus* populations compared to those in South America and it has been suggested that this is related to the greater level of habitat fragmentation and localization of livestock which discourages bats dispersal, as may be the case on the island of Trinidad [73, 76].

Our results indicate comparatively high genetic diversity in Trinidad, which is surprising for an island and may be the result of periodic inward bat dispersal (gene flow) from South America, which so far has effectively off-set any loss of diversity due to founder effects or isolation by distance, and female philopatry. We found that 94% of genetic variance was due to intra-population variation, which is likely related to female philopatric behaviour [69] with restricted dispersal around natal roosts resulting in more distinct groupings. Mammalian cytoplasmic genomes are thought to be almost exclusively maternally inherited [80] and not expected to display genealogical patterns representative of the history of the population, especially where sex-biases are thought to affect dispersal behaviour [81]. Male-biased dispersal with female philopatry, as seen in this study, has previously been demonstrated for both New World [52, 74] and Old World [82, 84] bat populations with juvenile males contributing more to genetic variability on account of their greater dispersal behavior [13, 69]. For vampire bats, there have been reports of juvenile male vampires from satellite colonies moving up to 100 km [13].

The geographic range of *D. rotundus* spans Mexico and all of Central and South America, including Trinidad, the Venezuelan island of Margarita and several smaller islands along the coast of Peru, Chile and Brazil [14, 67, 85–87]. To date there is no evidence of northern movement of vampire bats along the Caribbean archipelago past Trinidad, even though other Caribbean islands may provide adequate habitats. This restricted island range is thought to be due to the historical absence of large indigenous neotropical mammalian fauna during the pre-Columbian era [88, 89], an artefact of Caribbean island biogeography, before the ubiquitous presence of domestic livestock, which in addition to the distance between islands may have hindered the movement of this bat species up the islands, in contrast to other (non-hematophagous) bat species. However, given the contemporary increasing vampire bat population in Trinidad [90] and concomitant decline in livestock populations [91], high competition for resources could prompt aberrant bat movement into nearby naïve Caribbean island territories such as Tobago (42 km) and less likely onwards to Grenada (145 km).

It has also been suggested that climate change could result in north-westerly expansion of the vampire bat continental range from Central America to include Texas and Louisiana in North America [92], with additional potential habitats in Florida, California and Arizona [93–96] where there is fossil evidence for the existence of *Desmodus* bats thousands of years ago [94, 96]. However, for populations to be established especially on the islands, female *D. rotundus* bats must also demonstrate long-range movement which appears to contradict their philopatric behaviour. The successful establishment of the Trinidadian *Desmodus* island populations despite the intense male-biased migration may then be related to the persistence of relic populations since the separation of the island from the mainland. Therefore, although favourable climatic and habitat conditions may exist to host vampire bat populations within naïve (island and continental) territories, intensely male biased dispersal may limit the establishment of new populations especially on island territories across sea boundaries. Consequently, our genetic findings predict that other islands in the Caribbean appear to be at low risk of invasion by *D. rotundus* bats and based on the relative isolation of the island population has further important implications for *Desmodus* population control activities within Trinidad.

The oversea migration rate estimated in the current study for the Trinidadian *Desmodus* bat (i.e. 3.3 migrants per year) is higher than the rate (0.2 migrants per year) seen for over land migration between highly structured clades within Brazil located at least 100 km apart [51]. The majority of Trinidadian bats identified as having recent migrant ancestry (based on their microsatellite composition), were from the south-western peninsula close to Venezuela (12 km) suggesting more frequent entry from this point. However, the numbers are small and inference is limited by the fact that these bats may not be first generation migrants. Mark and recapture studies may provide a more comprehensive understanding of this movement.

Previous studies have demonstrated RABV circulation or exposure amongst the northern bat population [91, 97–99] with bat and viral introductions via the north-western peninsula also suggested [11, 91, 100, 101], albeit less frequently than from the south-western peninsula. The apparent bias toward southern viral entry has been proposed due to the more frequent presentation of rabies cases in the south on account of the concentration of livestock prey in that region [91]. Our previous study of Trinidad RABVs identified two major bat RABV lineages (Trinidad I and II) that were largely temporally defined and geographically distinct [99]. Trinidad I included 12 sequences from 1997–2000 (all but two of which were from the northeast of the island) and Trinidad II comprised 24 sequences from 2007 – 2010 and one from 2000, all from the southwest of the island, that clustered into two sub-lineages (IIa and IIb). Phylogeographic analyses suggested that Trinidad I, IIa and IIb each arose from independent introductions from the South American mainland (1972, 1989 and 2004 respectively) which the results of the current study suggest to be via male-dominated bat movement.

The limited mainland to island *D. rotundus* bat movement suggested by our study is consistent with our previously published observation of RABV outbreaks in Trinidad arising from relatively infrequent RABV introductions from the mainland [28], and suggests that determinants of mainland to island bat movement may be important drivers of vampire bat-transmitted rabies outbreaks in Trinidad, with epizootic spread promoted by intra island bat dispersal. Vampire bat population control on the island of Trinidad is therefore a critical component of rabies prevention and control programs to proactively prevent the incursion and establishment of new RABV lineages. In addition to measures within the island, to effectively reduce risks for RABV translocations via vampire bats, targeted control at the highlighted entry points to discourage bat incursions may be useful. For example, the implementation of livestock-free zones within the flight range of *D. rotundus* (i.e. 8 km in the southwest peninsula and 19 km in the northwest peninsula) may deter the inward movement of vampire bats to the island from the mainland. However, this may not address RABV introductions by other species with different migration patterns and flight ranges. Similar studies on other bat species are therefore recommended. Furthermore, consideration should also be given to the fact that RABV infection may either decrease successful viral dispersal due to the death of rabid bats or increase viral dispersal by causing aberrant migration behaviour [102, 103].

The spatial distributions of Trinidadian RABV lineages relative [28] to the *Desmodus* subpopulations identified in the current study suggest that the RABV Trinidad I lineage may have been introduced to and initially spread amongst the DR2 bats (presumably via entry from the north-west peninsula), while Trinidad II lineages likely entered and were maintained by DR1 bats in the southwest peninsula. However, since the Trinidad RABV sequences were derived from livestock and not bats [28], they represent single transmission events and may not fully represent the epidemic spread of these lineages within the bat population. Also, given that bat population structure is thought to have an inverse relationship with the rate of pathogen invasion and positively influence viral persistence [104], further investigations are necessary to determine the effect that *Desmodus* population separation by female philopatry may have on the dynamics of bat-transmitted viruses such as rabies on the island.

Although islands located near continents are rarely noted to develop unique species and have limited endemic species due to low rates of evolution and speciation [24], our results showed that *D. rotundus* bats from Trinidad were significantly smaller than those from the mainland. Morphological variation (excluding gender dimorphism) of *D. rotundus* throughout its continental range is well documented, and although some studies have suggested the possibility of sub-speciation [73, 74], others have indicated that neither morphological nor genetic variation was significant enough to warrant subspecies status [47, 105]. To our knowledge, this is the first study to demonstrate morphological (body size) differences between island and mainland populations of *Desmodus* bats. Such variability between island and mainland vertebrates has been widely documented and attributed to insular adaptations related to the small gene pool and restricted gene flow [106]. Given the close proximity of study sites, the morphological variability demonstrated in this study cannot be attributed to climatic plasticity [105]. The smaller size of the Trinidadian *D. rotundus* may in fact be an example of insular dwarfism [107, 108] which may have been an evolutionary adaptation to the limited food resources (prey species) associated with island biomes [109]. Most terrestrial animals currently found on this island are smaller in size than their continental counterparts because the natural environment does not favor the evolution of large terrestrial mammals [110]. A similar selective pressure was thought to have been instrumental in the extinction of the larger species of *Desmodus* from both within (e.g., *D. draculae* – Belize [111], Brazil [112], and Venezuela [113] and Mexico [114]) and outside (e.g., *D. archaeodaptes* – Florida [113] and *D. stocki* – Arizona [115], West Virginia [116], Florida [117], from North America and *D. putajudensis* from Cuba in the Greater Antilles [118–119]) of their current endemic range as a consequence of the extinction of Late Quaternary megafauna [94, 119].

Apart from environmental adaptation, *Desmodus* morphological diversity could also have been the result of genetic drift and a stepping-stone pattern of gene flow [105]. However, a more thorough genetic analysis including more comprehensive sampling would be necessary to make this distinction. In addition, future studies may exclude bats that recently fed and proportionally sample male and female bats to limit the introduction of potential biases.

## Conclusions

We have demonstrated that *D. rotundus* bats in Trinidad are genetically and morphologically distinct from their counterparts on the nearby South American mainland despite a degree of unidirectional gene flow from the mainland to Trinidad via male-biased migration (estimated at 3 bats per generation). Within Trinidad, two spatially segregated and genetically distinct *D. rotundus* subpopulations (DR1 and DR2) were identified at the mitochondrial level which the data suggests arose from different regions of South America. However, nuclear homogeneity of the Trinidad *Desmodus* population suggests male dominated intra-island movement facilitates gene flow between Trinidadian mitochondrial subpopulations, which may also be responsible for intra-island viral translocation. Further investigations into the relationship between these *D. rotundus* subpopulations and Trinidadian RABV lineages will further contribute to the development of contemporary evidence-based animal health and public health policies. These include more efficient vampire bat population control activities and rabies prevention programs in Trinidad, which protects both humans and livestock without subjecting other species of bats and their ecosystems to unnecessary and counterproductive persecution stemming from fear and misinformation.

## Acknowledgements

We are grateful to the numerous institutions and individuals that assisted with the acquisition of bat tissue samples from museum voucher specimens: Nancy Simmons and Eilenne Westwig (American Museum of Natural History), Robert Bradley and Heath Garner (Texas Tech University), Maria Eifler (Kansas University, Natural History Museum & Biodiversity Institute); Burton Lim (Royal Ontario Museum), Francois Catzeflis (University of Montpellier), Anne Lavergne and Benoit de Thoisy (Institut Pasteur de la Guyane) and Mike Rutherford (The University of the West Indies Zoology Museum). We thank the Anti-Rabies Unit and the Wildlife Section of the Ministry of Agriculture, Land and Fisheries, Trinidad and Tobago and the Trinidad and Tobago Bat Conservation and Research Unit for support in the field in Trinidad, and personnel from the Ministry of Agriculture, Animal Husbandry and Fisheries, Suriname and the Guyana Livestock Development Authority, Guyana for field assistance. We acknowledge the Department of Geomatics, Engineering and Land Management, The University of the West Indies, St. Augustine Campus and the Geographic Information Systems Unit, Ministry of Agriculture, Land and Fisheries for the provision of digital data layers for the maps.

## Supporting Information

**S1 Table: Details for *D. rotundus* specimens (n=194) used for microsatellite genetic analysis**

[Country, location and year collected, sex and source].

**S2 Table: Details for bat *cyt b* sequences used phylogenetic analyses** [Country, location and year that bat from which sequence was derived was collected, data set, Genbank accession number and phylogenetic placement (clade DR1 or DR2)].

**S1 Figure: Description of bat dispersal models tested for bat movement between Trinidadian clades and the mainland and within Trinidadian clades**.

**S3 Table: Summary of percentage nucleotide identities between and within country *D. rotundus* populations based on the *cyt-b* gene.**

**S4 Table: Number of alleles per loci by sample population.**

**S5 Table: Summary statistics for null allele frequencies (*r*) per locus by sampled population**

**S2 Figure. ML tree based on *D. rotundus cyt b* sequences (*n*=154; 382bp) from Trinidad and mainland South American countries together with corresponding sequences from *Diphylla ecaudata*.**

**S3 Figure: Number of observed and expected homozygotes (Hobs and Hexp) at each locus for all country populations.**

**S4 Figure: Graphical representation of the estimated probability of data for each k value (k=1-8) as determined by STRUCTURE analysis.**

**S5 Figure. Map illustrating the number of monitored bat roosts against the number of bovine farms and location of forested areas.**

## Data availability

Sequence data are available on Genbank (see **S2 Table** for accession numbers). Microsatellite data available from authors upon request.

